# Immediate early genes act in the medial hypothalamus to promote adaptation to social defeat

**DOI:** 10.1101/2025.09.09.675209

**Authors:** Rahul Sureka, Silvia Natale, Sukrita Deb, Demetra Avincola, Cornelius T. Gross

**Affiliations:** European Molecular Biology Laboratory (EMBL), Neurobiology and Epigenetics Unit, EMBL Rome, Via Ramarini 32, 00015 Monterotondo, Italy; Department of Bioengineering, University of Pennsylvania, 415 Curie Boulevard, Philadelphia 19104, USA; Department of Molecular and Cellular Physiology, Stanford University, 279 Campus Drive, Stanford, California 94305, USA; Sapienza University, Masters Programme in Neurobiology, Piazzale Aldo Moro 1, 00185 Roma, Italy

**Keywords:** Immediate Early Genes (IEG), Social fear, Innate fear, MK801

## Abstract

Territorial animals must moderate their social aggression and avoidance behaviors in a manner that maximizes their access to resources and fitness. The ventrolateral division of the ventromedial hypothalamic nucleus (VMHvl) has been shown to control both social aggression and avoidance in mice, and emerging data show that neural plasticity within VMHvl can drive the experience-dependent adaptation of these behaviors. Here, we investigated the contribution of immediate early gene (IEG) function in supporting this plasticity. In initial experiments, we found that downregulation of the IEG cFos in VMHvl did not significantly moderate the long-term increase in social avoidance seen following an experience of social defeat. However, local knockout of the IEG master regulator Serum Response Factor (SRF, *Srf*) was able to blunt the impact of social defeat on social avoidance and prevented the effect of social defeat on local optogenetic-evoked social avoidance behavior, demonstrating a critical role for subcortical IEG activity in social defeat-induced behavioral plasticity. To test whether NMDA receptor dependent plasticity might be involved in this effect, we locally infused the NMDA receptor antagonist MK-801 into VMHvl and assessed the impact of social defeat. Unexpectedly, MK-801 treatment led to an increase in social defeat-induced avoidance, pointing to the existence of opposing IEG and NMDA receptor-dependent adaptive mechanisms in medial hypothalamus. These findings suggest that multiple neural plasticity mechanisms are likely to be at play in the hypothalamic nuclei supporting innate behavior adaptation and show how IEG blockade can be used as a genetic tool to systematically map neural plasticity in the brain.

**Significance Statement:** The role of IEGs in experience-dependent plasticity has not been investigated in subcortical structures until now. Our findings suggest IEGs expression is essential for adaptive behavioral changes in the VMHvl following social defeat and can potentially be used to map behavioral adaptation across brain regions. Our study also indicates a potential use of IEGs as targets of gene therapy for mitigating maladaptive behavioral adaptations.

## Introduction

Transcription is widely appreciated to be necessary for the maintenance of long-term changes in synaptic plasticity. For example, high-frequency electrical stimulation in neuronal cultures leads to a rapid rise in excitatory synaptic strength that persists for hours after the stimulation has ceased. Pre-treatment with inhibitors of RNA polymerase has little effect on the initial rise in synaptic strength, but blocks the persistence of that increase over time, suggesting that the production of new gene products is required for the long-term consolidation of synapse potentiation and/or for the realization of other slow neural adaptation mechanisms (1–4). While a comprehensive list of the essential transcriptional cascades involved in supporting different types of synaptic and cellular plasticity is lacking, work has shown that a subclass of genes, called immediate early genes (IEGs), that are rapidly induced during high frequency electrical stimulation or learning-related behavioral experiences, are essential for the maintenance of excitatory synaptic plasticity (5, 6). Many of the most prominent IEGs (e.g., cFos) are transcription factors that subsequently bind to and activate a large number of target genes that are thought to be essential for supporting cellular plasticity (e.g., cytoskeleton, synaptic scaffolds, receptors, channels [7–9]). Work has shown that IEG activation depends on a rise in intracellular calcium and the activation of calcium-dependent signaling cascades. Blocking of critical calcium-dependent transcriptional activators such as cyclic-AMP response element binding protein (CREB) and serum response factor (SRF) using dominant negative strategies or knockdown has been shown to block IEG induction and abrogate activity-induced synaptic plasticity changes, providing an avenue for how synaptic activity can trigger IEG-dependent gene expression cascades (10–16).

While these experiments argue for a key role of IEGs in activity-dependent excitatory synaptic plasticity, few studies have demonstrated their role in experience-dependent synaptic plasticity *in vivo* or examined their contribution to other types of neural plasticity, such as changes in intrinsic properties, structural plasticity, or adaptive changes associated with non-neuronal cells (17–22). Moreover, work on IEGs in neural plasticity in mammals has so far been restricted to cortical structures, and little is known about the role of IEGs in behavioral adaptation mechanisms supported by subcortical innate behavior pathways, despite growing evidence for experience-dependent synaptic plasticity in these areas. Such work could be important because a major goal of current behavioral neuroscience research aims to map and understand the brain mechanisms that support adaptation to experience and how these might underlie maladaptive behavioral states. If IEGs were to play an essential role in neural plasticity across brain structures, they could be used as a tool to systematically map, manipulate, and understand the mechanisms underlying behavioral adaptation.

Here we sought to examine the role of IEGs in the medial hypothalamus on the ability of mice to adapt to social adversity. It is well established that in a wide variety of mammals, exposure to a short experience of social defeat at the hands of a dominant animal is associated with a persistent reluctance to interact with other animals, a form of generalized social anxiety. We used a combination of shRNA and conditional knockout strategies to block IEG expression in the medial hypothalamus, a brain network required for the expression of instinctive social threat responses, and examine their contribution to the behavioral adaptation to social defeat. We discovered an essential role for IEGs in subcortical pathways in this form of behavioral adaptation, and gathered evidence for the presence of multiple, functionally antagonistic plasticity mechanisms in these structures. Our findings suggest that IEG blockade can be used to systematically map experience-dependent neural plasticity at the cell-type level across the mammalian brain, a major goal of current behavioral neuroscience research.

## Results

### cFos induction in VMHvl is not required for behavioral adaptation to social defeat

To confirm previous observations that documented the induction of IEGs in the VMHvl following social defeat (23–25) we subjected adult male mice to social defeat by an aggressive male intruder mouse for a period sufficient to elicit social defeat (6-10 biting attacks) of the resident, returned the mouse to its home cage for 90 minutes and then euthanized the animal for immunofluorescent detection and quantification of cFos+ and Egr1+ cell density in the VMHvl (**Figure 1ab, Figure S1**). As expected, animals that underwent social defeat showed significantly higher density of both cFos+ (Student’s t-test, T = -5.89, *P* = 0.00036) and Egr1+ (Student’s t-test, T = -25.8, *P* = 5.4×10^-9^) cells in the VMHvl compared to non-defeated controls, confirming previous reports (26, 23, 25, 24).

**Figure 1.**
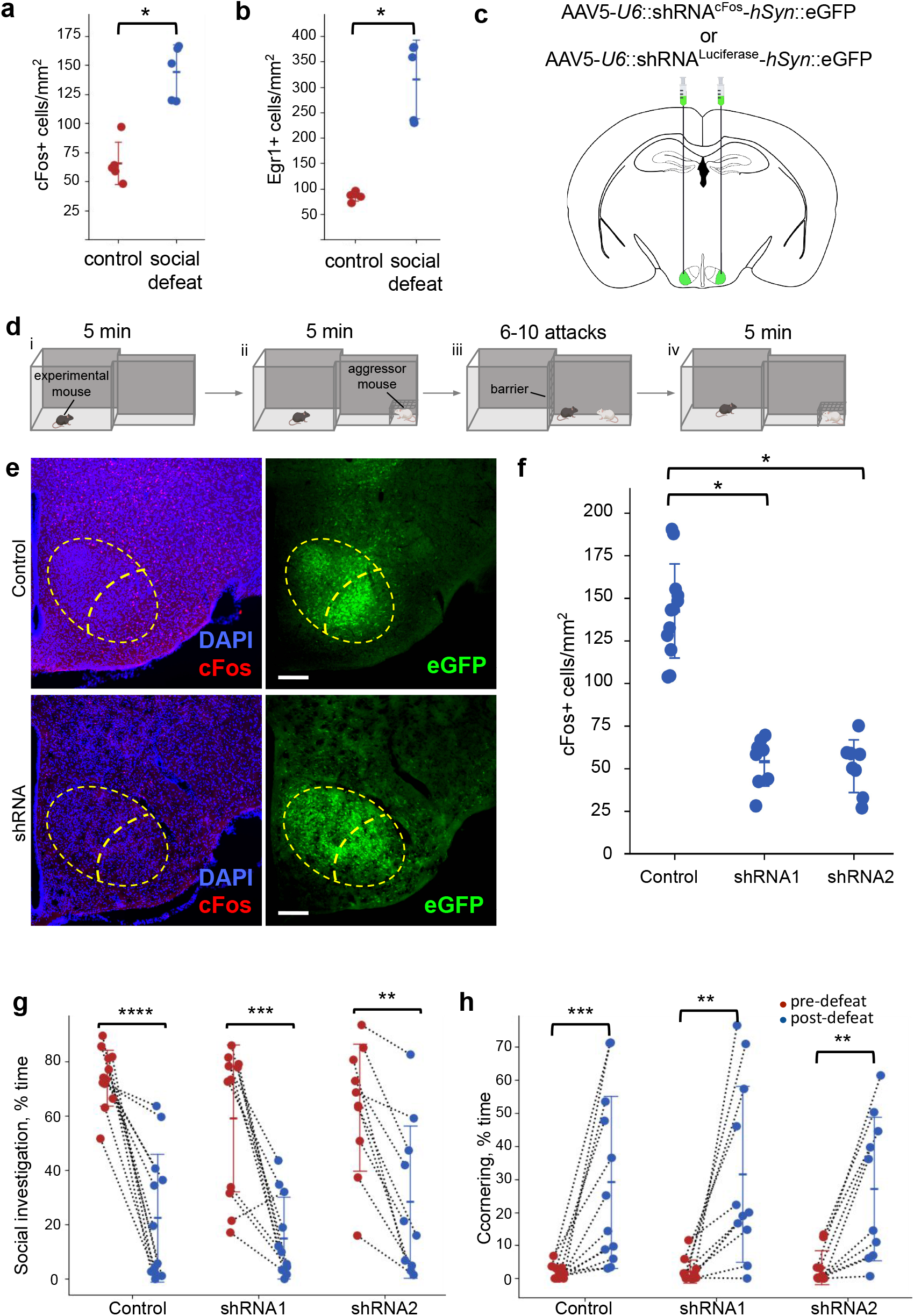
cFos knockdown in VMHvl does not affect social defeat responses in mice. Quantification of IEG expression in VMHvl 90 minutes after social defeat reveals a significant increase in **(a)** cFos (N = 5, *P* = 3.6e^-4^) and **(b)** Egr1 (N = 5, *P* = 5.2e^-9^) levels compared to control animals. **(c)** Anti-c*Fos* or control shRNA were expressed bilaterally in VMHvl by local stereotactic viral-mediated transfection. **(d)** The test mouse was habituated to the experimental apparatus (comprising a large safe chamber and a narrow corridor) for 10 minutes every day for 1 week. On the defeat day, the mouse was (**i**) habituated to the apparatus for 5 minutes, **(ii)** allowed to freely explore the apparatus with a conspecific male mouse restricted to a wire mesh cage at the far end of the corridor for another 5 minutes, **(iii)** restricted to the corridor with the wire mesh cage removed and subjected to social defeat for 6-10 attacks, and **(iv)** again allowed to freely explore the apparatus for 5 minutes. The three phases were repeated the next day and animals were sacrificed for immunostaining 90 minutes after defeat. Behavior from phase (ii) on the pre-defeat and post-defeat days was compared as a measure of the impact of social defeat on social avoidance. **(e)** Representative images of cFos expression overlaid with DAPI in control and cFos knockdown animals (left top and bottom, respectively) and spread of viral expression carrying control or shRNA vector (right top and bottom, respectively) in the same animals (scale bar = 100 µm). **(f)** Quantification of cFos expression in VMHvl of control and shRNA knockdown animals after social defeat (N = 8-12, *Post hoc* Tukey’s: shRNA1: *P* = 0.00, shRNA2 = 0.00). **(g)** Quantification of social investigation pre-defeat (red) vs. post-defeat (blue) in control and cFos knockdown animals (N = 8-12, *Post hoc* Tukey’s: pre-defeat vs post-defeat: control: *P* < 0.1e^-4^, shRNA1: *P* = 0.2e^-3^, shRNA2: *P* = 0.95e^-2^). **(h)** Quantification of cornering behavior pre-defeat (red) vs. post-defeat (blue) in control and cFos knockdown animals (N = 8-12, Wilcoxon Signed-Rank: Control: *P* = 0.4e^-3^, shRNA1: *P* = 0.19e^-2^, shRNA2: *P* = 0.19e^-2^).

Next, we tested whether the behavioral adaptations observed following social defeat were dependent on cFos expression in VMHvl. Expression of cFos was inhibited by local delivery of adeno-associated viruses (AAV) engineered to express one of two variants of a short hairpin RNA (shRNA) targeting two independent locations (exon 3 and 3’ UTR) in the cFos mRNA (shRNA1 vs shRNA2; **Figure 1c, Table 1**). Mice infected locally in VMHvl with AAV-shRNA1^cFos^-GFP, AAV-shRNA2^cFos^-GFP, or a control virus expressing a similar shRNA against the firefly Luciferase gene (AAV-shRNA^Luciferase^-GFP, not present in mice; **Figure 1c**) were subjected to social defeat in a modified resident intruder assay (**Figure 1d**). On defeat day (Day 1) the experimental animals were first allowed to habituate to a chamber attached to a short corridor for five minutes, followed by exploration for a further five minutes in the presence of an aggressive male intruder mouse restricted to a wire mesh cage placed at the far end of the corridor, during which period the fraction of total time spent investigating the intruder (% social investigation) as well as the fraction of time spent in the far corners of the main chamber (% cornering) were recorded. Subsequently, the barrier between the chamber and corridor was closed to confine the experimental animal in the corridor and the wire mesh cage was removed to allow the intruder to defeat the resident mouse (max 6-10 attacks or 10 min). Following social defeat the intruder was again confined in the wire mesh cage, the barrier was removed, and the fraction of time spent investigating the intruder, as well as the fraction of time spent in the far corners of the main chamber, were recorded for five minutes. Mice were transferred back to their home cage. 24 hours later, the same procedure was repeated. The experimental mice were euthanized 90 minutes after defeat for histological assessment of cFos and GFP expression to confirm that the virus was expressed in the VMHvl (**Figure 1e**) and that this was associated with a significant reduction in cFos+ cell density in VMHvl when compared to animals infected with the control viruses (ANOVA – main effect of treatment: F[1, 29] = 59.8, *P* = 2.93×10^-10^; **Figure 1f**).

**Table 1.**
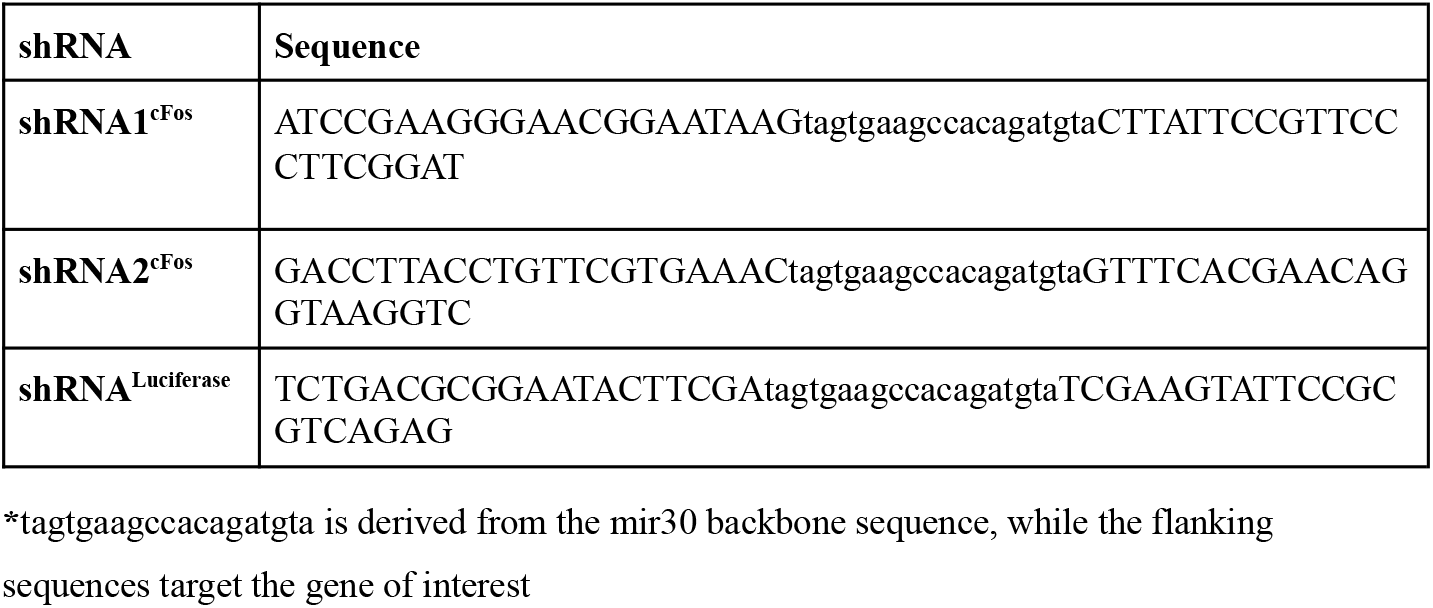
shRNA sequences

As expected, social defeat was associated with a significant reduction in time spent investigating the intruder (ANOVA – main effect of day: F[1,29] = 66.5, *P* = 2.67×10^-11^; **Figure 1g**). However, no significant effect of cFos shRNA treatment on social defeat-induced avoidance was detected for time spent investigating (ANOVA – treatment x day interaction: F[2,29] = 0.797, *P* = 0.455). Moreover, we also failed to detect any significant effect of cFos shRNA treatment on post-defeat cornering behavior (Kruskal-Wallis – effect of treatment: H[2] = 0.29, *P* = 0.86; **Figure 1h**), confirming that local knock-down of cFos in VMHvl does not impede the behavioral adaptation to social defeat.

### IEG induction in VMHvl is necessary for normal behavioral adaptation to social defeat

One explanation for our findings is that IEGs other than cFos might be involved in social defeat-induced behavioral plasticity (27). To downregulate multiple IEGs at once, we used a conditional knockout strategy to block expression of the IEG master regulator, Serum Response Factor (SRF, *Srf*; 28–30) locally in VMHvl. Mice homozygous for the Cre-conditional allele of *Srf* (*Srf*^cKO^) were injected into the VMHvl with an adeno-associated virus expressing either Cre recombinase (AAV-Cre-GFP) or a control vector (AAV-GFP) (**Figure 2a**) and subjected to social defeat testing (**Figure 1d**). Successful knockout of *Srf* was evaluated by quantifying the social defeat-induction of IEGs cFos and Egr1 in VMHvl (**Figure 2d**). Quantification of the density of cFos+ and Egr1+ cells in the area infected by the virus confirmed a significant downregulation of social defeat-induced cFos (Students t-test, T = 6.95, *P* = 3.29×10^-06^; **Figure 2b**) and Egr1 (Students t-test, T = 13.2, *P* = 4.81×10^-10^; **Figure 2c**) expression in *Srf*^cKO^ animals that received the Cre virus compared to controls, demonstrating a significantly reduced induction of IEGs in these animals.

**Figure 2.**
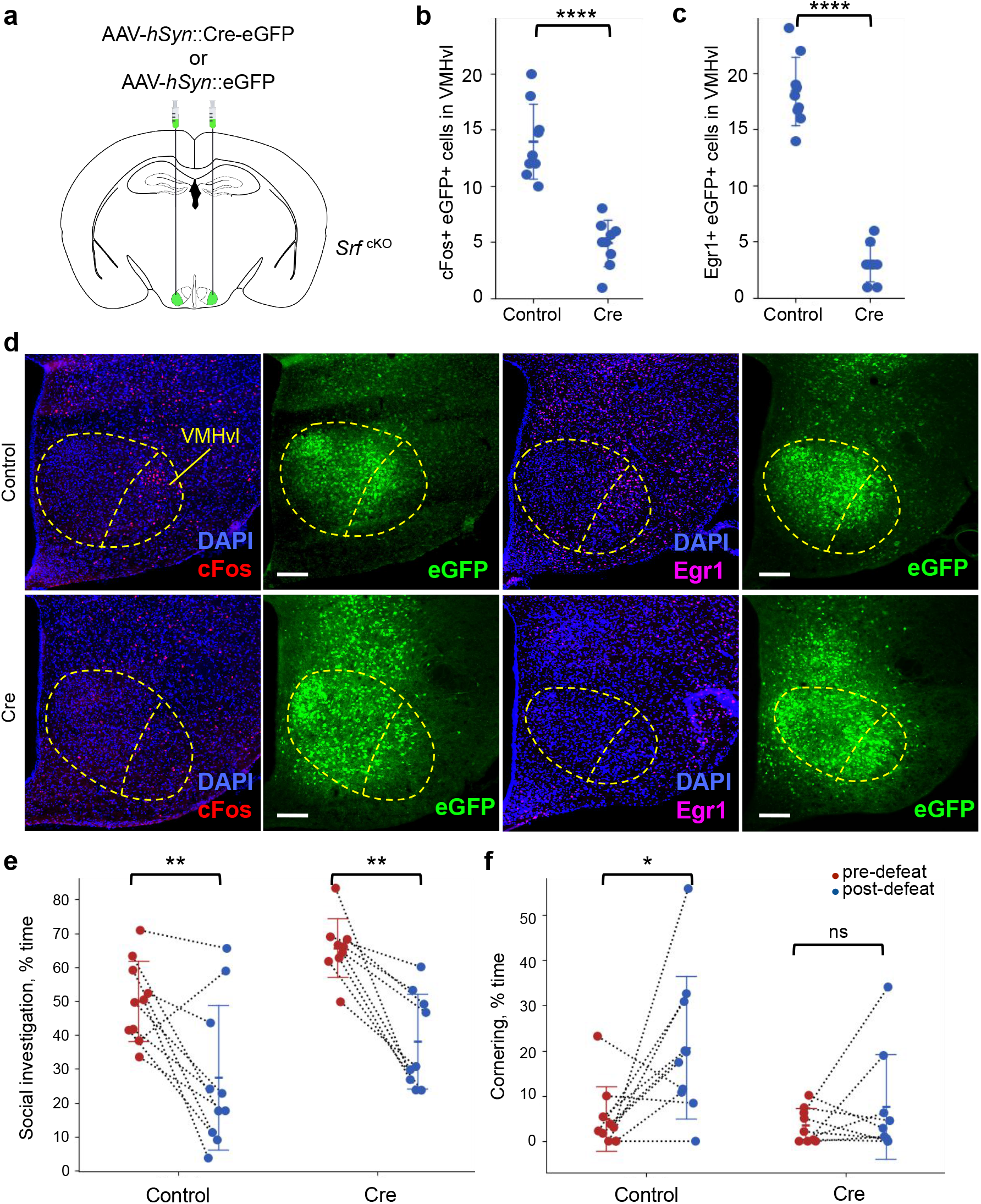
Local knockout of *Srf* in VMHvl partially blocks the effect of social defeat. **(a)** Cre recombinase or empty-vector control were expressed bilaterally in VMHvl by local stereotactic viral-mediated transfection in *Srf*^*cKO*^ animals. The animals were subjected to the same behavior testing as in Figure 1d. Quantification of co-expression of IEGs **(b)** cFos+ or **(c)** Egr1+ and Gfp+ nuclei in the VMHvl of control and Srf knockdown animals after social defeat (N = 9, *P* = 3.3e^-6^, *P* = 4.8e^-10^). **(d)** Representative images of GFP and (left) cFos or (right) Egr1 co-expression overlaid with DAPI staining in control and Srf knockdown animals (scale bar = 100 µm). **(e-f)** Quantification of social investigation and cornering behavior pre-defeat (red) vs. post-defeat (blue) in control and *Srf* knockout animals(Social Investigation: N = 9, *Post hoc* Tukey’s: Control pre-defeat vs post-defeat: *P* = 0.89e^-2^, Cre pre-defeat vs post-defeat: *P* = 0.22e^-2^, Cornering: N = 9, Wilcoxon Signed-Rank: Control: *P* = 0.028, Cre: *P* = 0.82).

However, despite a significant inhibition of IEG induction, both *Srf*^cKO^ mice treated with Cre-expressing or control virus showed a significant reduction in social investigation following social defeat (ANOVA – main effect of day: F[1,18] = 26.8, *P* = 1.0×10^-5^; **Figure 2e**) and no significant difference in response to social defeat was seen in Cre-expressing virus infected animals when compared to controls for social interaction behavior (ANOVA – treatment x day interaction: F[1,18] = 0.257, *P* = 0.615; **Figure 2e**). However, cornering behavior did reveal a significant effect of social defeat in control animals, but no significant effect in *Srf*^cKO^ mice. Although the non-parametric distribution of the data did not allow a direct assessment of the statistical interaction between social defeat and treatment for this measure, a pairwise comparison of post-defeat behavior between *Srf* knockout and control groups revealed significantly less cornering in the knockout group (Mann-Whitney U Test [post-defeat]: U = 70.0, *P* = 0.045) corroborating that observation that *Srf*^cKO^ animals were less susceptible to the effects of social defeat than controls. Together, these findings point to a role for IEGs in VMHvl in mediating the behavioral adaptation to social defeat, but in a manner that may be occluded by plasticity in other brain areas.

### IEG induction in VMHvl is required for evoked behavioral adaptation to social defeat

To more directly interrogate whether plasticity in the VMHvl pathway contributes to the behavioral adaptation to social defeat we turned to a method that depends on direct optogenetic activation of VMHvl to elicit social avoidance behavior. We and others (24, 25, 31) have shown that optogenetic activation of VMHvl can elicit short-latency social avoidance in mice and that this response is enhanced after social defeat (25, 24). Groups of homozygous male *Srf*^cKO^ and wild-type mice were infected in VMHvl with a cocktail of Cre-expressing virus and a virus expressing either a Cre-dependent channelrhodopsin (AAV-DIO-ChR2-YFP) or a control virus (AAV-DIO-YFP; **Figure 3a**) and implanted bilaterally with optic fibers centered over the VMHvl (**Figure 3b**). Mice were subjected to a modified social avoidance apparatus (24) **(Figure 3c**) in which a subordinate male was introduced into the home cage of the experimental mouse and following 30-60 seconds of social interaction, short pulses (20 Hz pulses for 30 s, ITI 60 s) of blue light were delivered to elicit behavioral responses. One day later, the experimental animal was subjected to social defeat in a novel cage by an aggressive male mouse. On the third day, the mouse was again tested for light-evoked social behavior responses in the same manner as before.

**Figure 3.**
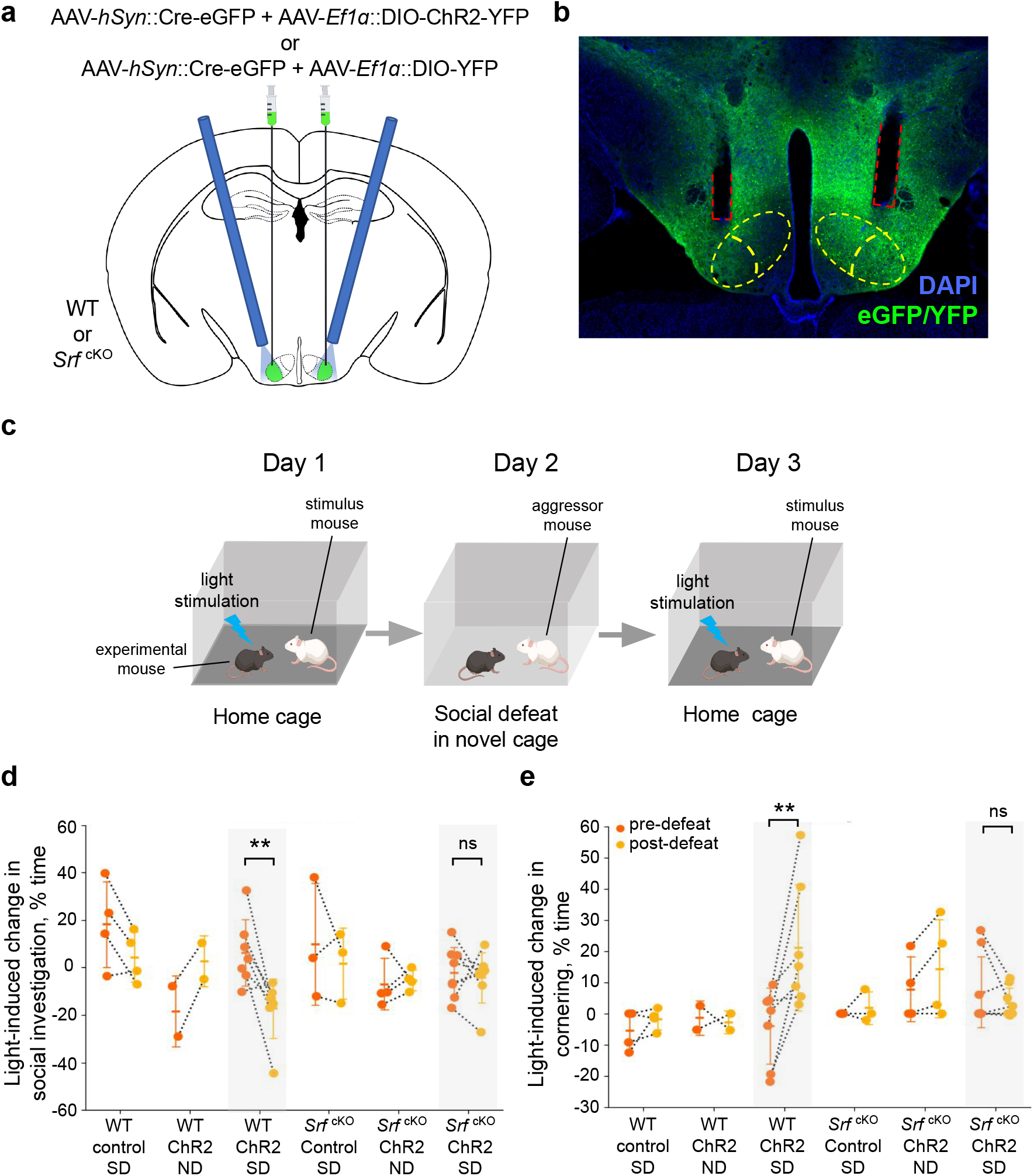
Local knockout of *Srf* in VMHvl blocks the effect of social defeat on optogenetics-evoked avoidance behavior. **(a)** Cre recombinase and either channelrhodopsin (ChR2) or empty-vector control were expressed bilaterally in VMHvl by local stereotactic viral-mediated transfection in WT or *Srf*^*cKO*^ animals in which bilateral optic fibers were implanted over VMHvl. **(b)** Representative image showing the transfection site of the virus and the placement of optic fibers (red dashed line). **(c)** Animals were tested in a modified social defeat testing protocol. Following one week of habituation, on the first testing day a subordinate male mouse was introduced into the home cage of the experimental animal and the experimental mice was stimulated 3-5 times for 30 seconds (ON, 20 Hz, 20 ms light pulse) interspersed by 30 seconds (OFF). On the second testing day, the experimental mouse was placed into a novel cage either together with an aggressive conspecific male mouse and subjected to social defeat for 6-10 attacks (social defeat, SD) or into an empty novel cage for 10 minutes (no defeat, ND). On the third testing day a novel subordinate male mouse was introduced into the home cage of the experimental mouse and the experimental mouse was stimulated 3-5 times as on Day 1. **(d)** Quantification of the light-evoked social investigation and cornering behavior (average time spent during ON minus OFF) pre-defeat (Day 1, orange) vs post-defeat (Day 2, yellow) in WT and *Srf*^*cKO*^ animals (N = 2-7, *Post hoc* Tukey’s: Social Investigation: WT ChR2 SD pre-defeat vs post-defeat: *P =* 0.53e^-2^, *Srf*^cKO^ ChR2 SD pre-defeat vs post-defeat: *P =* 0.98, Cornering: WT ChR2 SD pre-defeat vs post-defeat: *P =* 0.65e^-2^, *Srf*^cKO^ ChR2 SD pre-defeat vs post-defeat: *P =* 0.95)

Following social defeat, optogenetic stimulation of VMHvl neurons elicited a greater reduction in social interaction toward the subordinate mouse when compared to light stimulation on the day before social defeat, and this difference was significantly reversed in *Srf*^cKO^ mice receiving the Cre-expressing virus (social interaction: ANOVA – treatment x day interaction: F[1, 13] = 5.99, *P* = 0.021; cornering: ANOVA – treatment x day interaction: F[1, 13] = 9.015, *P* = 0.0058; **Figure 3d** and **Figure S2**). On the other hand, no effect of interaction between treatment and day of testing on light-evoked responses was seen in control animals that either lacked ChR2 expression, were not subjected to social defeat, or who carried the wild-type allele of *Srf* (WT; **Figure 3e**). To confirm interactions between social defeat and *Srf*^cKO^ genotype for light-evoked responses we also carried out a comprehensive statistical assessment across multiple behavior measures (social investigation, cornering, immobility, and digging). This analysis revealed a significant interaction between *Srf*^cKO^ genotype and day of testing for all four behavior measures (MANOVA – genotype x day interaction: λ = 0.165, F = 23.2, *P* = 0.00) and argues for a robust impact of *Srf* knockout on local social defeat-associated neural function changes.

### Local delivery of MK801 enhances behavioral adaptation to social defeat

Previous studies have shown that IEG activation is required for long-term excitatory synapse potentiation (32, 29, 33). However, experience-dependent brain plasticity is known to involve a panoply of functional changes that include NMDA receptor-dependent changes in excitatory synaptic strength, non-NMDA receptor-dependent changes in synaptic function, intrinsic neural properties, functional connectivity, and glia and vascular function. As a first step toward understanding which, if any, of these plasticity mechanisms might be involved in social defeat-associated behavioral adaptation, we delivered MK801 locally to VMHvl to block NMDA receptor-dependent plasticity (**Figure 4a**). Mice were implanted with guide cannulae positioned over the VMHvl (**Figure S3**) and injected with either a low or high dose of MK801 twenty minutes before being subjected to social interaction testing following a procedure similar to that used above for examining the impact of IEG knockdown but where the wire mesh barrier was located at the border between the chamber and corridor (**Figure 4b**). Behavioral adaptation to social defeat was evaluated 24 hours later by comparing social interaction measures before (day 1) and after defeat (day 2). A significant dose-dependent interaction between MK801 treatment and social defeat emerged (social avoidance, exploration: MANOVA – treatment x day: λ = 0.38, F = 8.10, *P* = 0.002). Unexpectedly, MK801 treatment was associated with a dose-dependent enhancement of social avoidance and behavioral inhibition seen after social defeat (ANOVA - main effect of treatment: Avoidance: F[2,38] = 3.337, *P* = 0.047, Exploration: F[2,38] = 2.39, *P* = 0.107, main effect of defeat: Avoidance F[2,38] = 18.747, *P* = 0.125e^-3^, Exploration: F[2,38] = 23.263, *P* = 0.29e^-4^) (**Figure 4c,d**). These findings support a role for local NMDA receptor-dependent mechanisms in antagonizing rather than promoting adaptation to social defeat, and when considered together with our findings in *Srf*^cKO^ mice, argue for the existence of parallel positive and negative adaptive mechanisms to social defeat.

**Figure 4.**
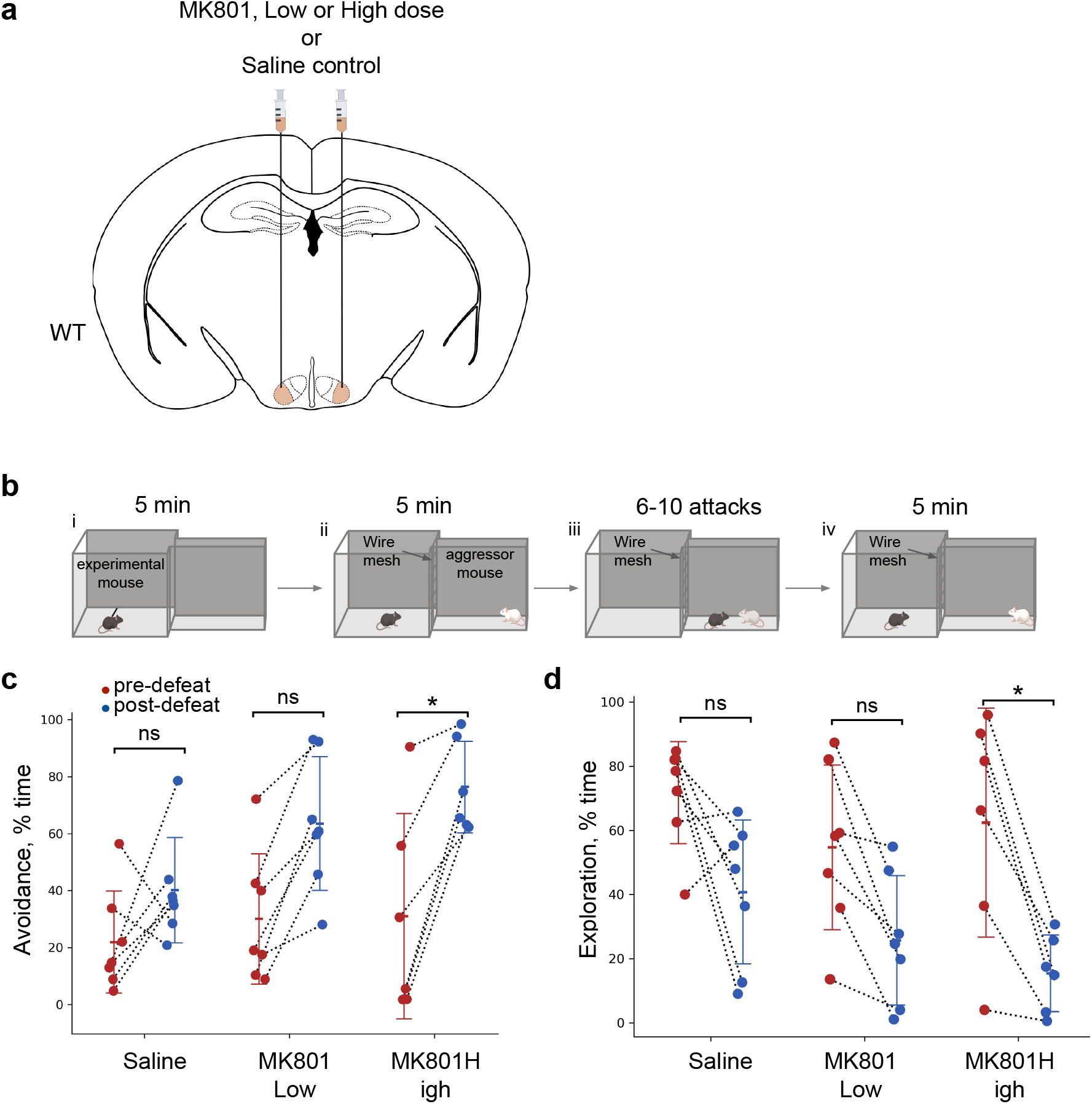
Local delivery of NMDAR antagonist MK801 in VMHvl enhances social defeat-induced avoidance behavior. **(a)** Guide cannulae were implanted bilaterally to deliver saline or either a low (0.025 ng) or high dose (0.050 ng) of MK801 locally into VMHvl. **(b)** The test mouse was habituated to the microinjector and experimental apparatus (comprising a large safe chamber and a narrow corridor) for 10 minutes every day for 1 week. On the defeat day (day 1), saline or MK801 was injected using an automatic microinjector pump 20 minutes before the animal was (**i**) habituated to the apparatus for 5 minutes, **(ii)** allowed to freely explore the chamber with a conspecific male mouse restricted to the corridor by a wire mesh barrier for another 5 minutes, **(iii)** restricted to the corridor with the aggressor and subjected to social defeat for 6-10 attacks, and **(iv)** again allowed to freely explore the chamber with the aggressor restricted to the corridor for 5 minutes. Phases (i) and (ii) were repeated 24 hours later (day 2) with a novel conspecific. Quantification of **(c)** social avoidance and **(d)** exploration behavior pre-defeat (red) vs. post-defeat (blue) in mice injected with saline, low, or high doses of MK801 (N = 6-7, Post hoc Tukey’s: Avoidance: Saline: *P =* 0.684, Low MK801: *P* = 0.1, High MK801: *P* = 0.02, Exploration: Saline: *P =* 0.147, Low MK801: *P* = 0.199, High MK801: *P* = 0.014).

## Discussion

We have applied virus-delivered shRNA and Cre-conditional genetic knockout tools to block the induction of IEGs in the mouse medial hypothalamus and examine their role in social defeat-induced social avoidance behavior. We found that IEG induction blockade partially reduced defeat-induced social avoidance, but fully blocked the increase in local optogenetic-invoked social avoidance seen after social defeat. These findings confirm the critical role of IEGs in behavioral adaptation and extend earlier findings of their role in cortical structures required for associative learning to subcortical structures required for innate stimulus adaptation.

In parallel, we investigated the impact of local NMDA receptor blockade on social defeat-induced social avoidance. We found that treatment with the NMDA receptor antagonist MK801 before social defeat was associated with an unexpected increase in social avoidance one day later. These findings suggest that NMDA receptor and IEG-dependent local plasticity mechanisms have opposing roles in the behavioral adaptation to social defeat. Previous studies have argued that IEG induction in subcortical structures may not depend on calcium entry via NMDA receptors (34) and thus may support independent transcriptional activation pathways. Similarly, signaling via CREB, a known downstream effector of NMDA receptor activation in cortical structures is not altered in *Srf* cKO mice (29).

Our observation that IEG blockade in VMHvl fully inhibited ChR2-evoked, but only partially countered baseline changes in social behavior following social defeat, suggests a functional redundancy of plasticity in multiple brain pathways in the adaptation to defeat. Indeed, plasticity in frontal cortical areas has been shown to be induced following social defeat (35–39) and either systemic or local delivery of NMDA receptor antagonists in the frontal cortex interferes with the impact of social defeat. We note that our observation that IEG blockade in VMHvl reduced the effect of social defeat on some measures of baseline behavior (**Figure 2f**), supports the argument for a masking of the expression of plasticity in medial hypothalamus by other brain pathways, rather than the alternative argument for independent mechanisms underlying baseline and ChR-evoked behavioral plasticity. This view is supported by the significant impact of local NMDA receptor blockade observed on baseline social avoidance activity. Nevertheless, the discrepancy between the baseline and ChR2-evoked responses of IEG blockade does raise the question as to the conditions under which the contribution of VMHvl to behavioral adaptation might be more pronounced. Finally, our finding that local NMDA receptor antagonism potentiates (**Figure 4cd**), while systemic or frontal cortical NMDA receptor blockade prevents (40–43) the impact of social defeat points to antagonistic contributions of similar plasticity mechanisms between cortical and subcortical brain pathways. This conclusion stands even if the tissue diffusion of MK801 could not be directly tracked in our experiments and may have extended to parts of the hypothalamus or thalamus outside of VMHvl (**Figure 4a**).

Although a comprehensive electrophysiological characterization of social defeat-induced neural plasticity changes in VMHvl is beyond the scope of the current study, our findings lead us to speculate that multiple IEG and NMDA receptor-dependent plasticity mechanisms are elicited in VMHvl during defeat. Previous studies have documented the potentiation of excitatory inputs to VMHvl during repeated experiences of aggression and complex changes in both excitatory and inhibitory synaptic currents in VMHvl following social defeat (44). Our work leads to the prediction that these mechanisms could be dissected by examining their differential dependence on IEGs or NMDA receptors.

In summary, we have found that IEG blockade can be used as a tool to block behavioral adaptation arising from neural plasticity in subcortical brain structures and used this approach to provide evidence for the recruitment of competing experience-dependent neural plasticity mechanisms. Work will need to be done to identify the sub-types of neural plasticity that depend on IEGs and compare and contrast them with those dependent on NMDA receptors or other, better understood mechanisms. Nevertheless, our findings show how such tools could be used to systematically carry out cell-type specific mapping of neural plasticity across the mammalian brain and help move the focus of current behavioral neuroscience from the study of neural activity to neural plasticity.

## Material and Methods

### Animals

All experimental procedures involving the use of animals were carried out in accordance with EU Directive 2010/63/EU with approval from the EMBL IACUC and the Italian Ministry of Health Licenses 541/2015-PR and 183/2024-PR. All non-transgenic animals (C57BL/6N mice) were bred locally as part of EMBL internal colonies. The *Srf*^cKO^ line (RRID: IMSR_JAX:006658) was a gift from Dr. Leszek Kaczmarek at the Nencki Institute, Warsaw, Poland. These animals were rederived in the C57BL/6N background and backcrossed to C57BL/6N for 10 generations. Thereafter, all experimental animals were derived from homozygous colonies maintained locally. Aggressor mice were singly housed CD1 adult retired male breeders (Charles River) screened for robust aggression (Franklin et al., 2017). Subordinate mice were 8-12 week-old BALB/c males derived from internal EMBL colonies. All animals were maintained in a temperature and humidity-controlled environment with 12 hr light and 12 hr dark cycle, and food and water were provided *ad libitum* with biweekly changes of cages. One week before the initiation of the experiment, the experimental male mice were singly housed.

### Stereotactic surgeries

Mice were anesthetized with 3.5-4% isoflurane (Provet) in oxygen for 2-3 minutes and secured in a stereotactic frame (RWD). Anesthesia was maintained with continuous 1-1.5% isoflurane administration in oxygen through the nose cone. Body temperature was maintained at 37℃ with a heating pad, and the eyes were protected with a lubricant. The scalp was shaved and disinfected with Betadine. The skull was exposed, aligned to the stereotactic frame, and cleaned with 0.3% hydrogen peroxide solution. For cFos knockdown experiments, 0.2-0.3 µl (titer > 10^12^) AAV5-*U6*::shRNA1^cFos^-*hSyn*::eGFP, AAV5-U6::shRNA2cFos-hSyn::eGFP, or AAV5-*U6*::shRNA^Luciferase^-*hSyn*::eGFP virus was infused bilaterally into VMHvl (from Bregma L: +/-0.69, A/P: 0.97, D/V: 5.85) of 8-9 weeks old C57BL/6N mice. For Srf knockdown experiments, AAV5-*hSyn*::Cre-eGFP or AAV5-*hSyn*::eGFP virus was similarly infused into the VMHvl of *Srf*^cKO^ mice. For optogenetic experiments, AAV5-*hSyn*::Cre-eGFP + AAV5-*Ef1a*::DIO-ChR2 or AAV5-*hSyn*::Cre-eGFP + AAV5-*Ef1a*::DIO-YFP virus was bilaterally infused into the VMHvl (from Bregma L: -0.69, A/P: 0.97, D/V: 5.85 and L: +1.69, A/P: 0.97, D/V: 5.87 at 10^°^ angle) of *Srf*^cKO^ or C57BL/6N mice (WT) at the same coordinates. Five to ten minutes later, the capillary was slowly withdrawn, and optic fibers (0.22 NA, 200 mm core and 1.25 mm ceramic ferrule; RWD) were implanted (from Bregma L: -0.69, A/P: 0.97, D/V: 5.65 and L: +1.69, A/P: 0.97, D/V: 5.67 at 10^°^ angle). For NMDAR antagonist experiments cannulae were implanted bilaterally in C57BL/6N mice above VMHvl (from Bregma L: -0.69, A/P: 0.97, D/V: 5.65 and L: +1.69, A/P: 0.97, D/V: 5.67 at 10^°^ angle). All implants were secured to the skull using dental cement (Duralay). The wound was cleaned, and the skin was stitched around the implant. At the end of the surgery, 5 mg/kg of carprofen dissolved in saline was injected subcutaneously and mice were placed into a heated cage for two hours. Once the mice were alert, they were moved to a new cage and maintained in isolation with drinking water containing paracetamol (Tachipirina) for one week for recovery. Thereafter, the mice were singly housed for three weeks with normal food and water before experimentation. All viruses were produced by the EMBL Genetic and Viral Engineering Facility.

### Behavior testing

For most experiments (**Figures 1, 2** and **4**) the experimental apparatus consisted of a chamber (25 x 25 x 25 cm, plexiglas) with a 6 cm wide opening in one of the walls serving as the entrance and a corridor (30 x 6 x 25 cm, plexiglas) with two open ends. The corridor could be connected to the chamber via one of the open ends and the other end closed using a sliding door. The entrance between the chamber and the corridor could be closed using a sliding door. For the optogenetic experiment (**Figure 3**) social exposure to a subordinate animal was done in the home cage of the experimental animal, while social defeat was done in a novel, but identical cage to the home cage. For all experiments except the optogenetics experiment, the experimental animals were habituated to the apparatus for 5 min followed by 5 min of free exploration with an empty wiremesh cage at the far end of the corridor for five days immediately preceding the defeat day. For MK801 experiments (**Figure 4**) the animals were handled and the cannula plugged to an automatic injector (Hamilton) for local infusion of MK801 every day for 10 minutes before placing them in the experimental apparatus for habituation to the apparatus. On experimental day 1 (defeat day), the experimental animals were allowed to explore the apparatus for 5 min. MK801 or saline was infused 10 minutes before introduction into the apparatus. After 5 min, an aggressive conspecific (CD1) was restricted at the far end of the corridor in the wire mesh cage while the experimental mouse was allowed to freely explore for another 5 min. For animals in the No Defeat (ND) group, the experiment was terminated at this point. Thereafter, the aggressive conspecific was released from the wire mesh cage, and the animals were restricted to the corridor to permit social defeat of the experimental animal up to 10 attacks by the conspecific or 10 min (whichever was reached earlier). Following social defeat, the conspecific was restricted to the far end of the corridor in a wire mesh cage while the experimental animal was allowed to roam freely for another 5 min. The procedure for experimental day 1 was repeated on experimental day 2, and the experimental animal was sacrificed by perfusion and fixed 1.5 hrs after the social defeat to allow for IEGs immunofluorescent imaging of brain slices. Care was taken to ensure that the experimental animal was never exposed to the same aggressive animal on days 1 and 2. The bedding in the apparatus was changed, and the cage was wiped sequentially with ethanol and water between every experimental animal.

For the optogenetic experiments, the experimental animals were habituated to the experimental setup for 5 min in an open-top home cage for 3 days. For the following 4 days of habituation, handling, and restriction by scruffing were gradually introduced, after which the animals were again allowed to explore the home cage without its cover in the experimental setting. For the next 7 days, they were further habituated to plugging the optic fibers and allowed to explore the cage without the cover for 5 min with the cable connected. On the first day of the experiment, the experimental mouse was similarly plugged and allowed to habituate for 5 min in the home cage. Thereafter, a docile conspecific (BALB/c) was introduced into the cage. After 30-60 seconds of free exploration, the experimental animal was stimulated with light (30s, 20Hz) every 60 seconds for 3 cycles of light ON and OFF with 5-6 mW/mm^2^ of laser power. On the next day, the animals were placed in a novel, but identical cage. After 5 min of free exploration, an aggressive CD1 male was introduced and social defeat was allowed to take place for 6-10 attacks or 10 min (whichever was reached earlier). On the third day, the same protocol as experimental day 1 was followed. Care was taken to ensure that the experimental animals were not re-exposed to the same BALB/c animal. ChR2 was optically stimulated using a 465 nm laser connected to a power supply (PSU-III-LED, Thorlabs) and attached to a manual rotatory joint with 1-meter high-performance patch cables (Thorlabs). Power at the patch cable terminal was measured before and after each experiment with a portable optical power meter (Thor Labs) and stimulation trains were generated using the Pulser software. Mice with misplaced viral injections or optic fibers were excluded from the study.

### Behavioral data acquisition and annotation

Behavior was recorded (25 fps, Pylon software) using a ceiling mounted camera (acA1300-60gmHIR, Basler) connected with GigE. Behavioral annotation was carried out manually frame by frame using Solomon Coder software. All videos were cropped using the windows default video editing tool to only include the period being scored. All identifying features about the treatment of the animal were removed from the video filenames and logged in a separate file to ensure blind scoring of behavior. For all experiments except optogenetics, the behavior of the experimental animal in the 5 min free exploration period with the restricted conspecific before social defeat was scored for both days 1 and 2. For the optogenetic experiments the behavior of the experimental animal in the home cage upon introduction of the BALB/c was scored on day 1 and 3 of the experiment. The following behaviors were scored: social investigation – actively sniffing, facing, or grooming the conspecific or biting the wire mesh cage restraining the conspecific; cornering – staying in the far corner of the apparatus with the body against the wall of the apparatus; immobility – no body movement; digging – overturning and displacing the bedding with the front paws; avoidance – actively turning away or staying far from the conspecific. Any animals with misplaced viral infections or optic fiber implants were excluded from the analysis.

### Data analysis

All behavior scores were converted to percentage values by dividing by the total time of the video clips and compiled in tab-separated files. For the cFos, Srf knockdown, and MK801 experiments two-way ANOVA and Tukey’s post-hoc testing were used. For measures that did not meet the assumptions of ANOVA, either the Kruskal-Wallis test or the Mann-Whitney U Test for pairwise comparisons was used. For the optogenetic and MK801 experiments, MANOVA with *post hoc* ANOVA was used. cFos, Egr1, and GFP-expressing cells were manually counted from at least two sections per animal. The area of VMHvl was annotated and the area was calculated using the built-in tools in Qupath. Student’s T-test or one-way ANOVA was used to evaluate significant differences. All statistical analyses and plots were generated using custom scripts in Python.

### Immunohistochemistry and microscopy

One-and-a-half hours after the last experimental time point all animals (except those for optogenetics) were deeply anesthetized with 0.01 ml/g of Avertin and transcardially perfused with cold PBS (137 mM Nacl, 2.7 mM KCl, 20 mM NaH_2_PO_4_, 1.8 mM K_2_HPO_4_) followed by cold 4% PFA in 0.1M PB solution (75.4 mM NaH_2_PO_4_, 24.5 mM Na_2_HPO_4_). The brain was post-fixed in 4% PFA at 4^°^C overnight and coronal sections were sliced using a cryotome for immunofluorescent staining (thickness 10 µm). The sections were stored in PBS (supplemented with 0.1% NaN_3_) at 4^°^C. Anti-cFos staining was done according to the protocol outlined in (Krzywkowski et. al, 2020). Briefly, sections were washed with PBS, permeabilized with 0.2% Triton-X in PBS (PBST), blocked with 5% donkey serum in PBST, and incubated with cFos primary antibody (1:1500) (SC-52G, Santa Cruz) in PBST overnight with gentle shaking. Sections were washed three times in PBS, incubated with AlexaFluor 647 conjugated anti-rabbit secondary antibody (1:10000, A21244, ThermoFischer) in PBS for 2 hrs at room temperature, and washed twice with PBS. They were stained with DAPI (1:1000 in PBS) for 15 min, washed with PBS, and mounted on SuperFrost Plus slides (ThermoFischer) with Moviol. For Egr1 staining the same protocol as cFos was used except swapping the primary antibody with anti-Egr1 antibody (4154S, Cell Signaling) at a dilution of 1:1000. The brains of animals used for optogenetics were similarly processed as above one day after experimental day 3, and sectioned (thickness 20 µm) for checking implant localization and injection site.

## Acknowledgements

We thank Roberto Voci for expert animal husbandry and other members of the EMBL Laboratory Animal Resources (LAR) staff for their support. We thank the EMBL Genetic and Viral Engineering Facility (GAVEF, now GEVF) for viral production and the EMBL Light Imaging Facility (LIF) for microscopy support. This research was supported by an EMBL Rome interface grant to Mathieu Boulard and C.T.G., EMBL PhD Fellowship to S.D., Federico II University of Naples fellowship to S. N., European Research Council (ERC) Advanced Grant (AdG) TERRITORY #101097411 to C.T.G., and EMBL core funding. R.S. carried out all the experiments and analyzed and interpreted data, except the MK801 experiments that were carried out by S.N. with histology support from S.D. Initial experiments aimed at documenting evoked Egr1 expression were carried out by D.A. The research was conceived and manuscript written by R.S. and C.T.G with input from S.N.

## Supplementary Figures

***Figure S1***. Representative images of cFos and Egr1 expression overlaid with DAPI in social defeat (SD) vs. no defeat (ND) animals.

***Figure S2*. (a-b)** Quantification of immobility and digging behavior pre-defeat (orange) vs. post-defeat (yellow) in WT and *Srf*^*cKO*^ animals (N = 2-7, *Post hoc* Tukey’s: Immobility: WT ChR2 SD pre-defeat vs post-defeat: *P* = 0.519, *Srf*^cKO^ ChR2 SD pre-defeat vs post-defeat: *P* = 0.2e^-2^, Digging: WT ChR2 SD pre-defeat vs post-defeat: *P* = 0.997, *Srf*^cKO^ ChR2 SD pre-defeat vs post-defeat: *P* = 0.837)

***Figure S3***. Representative image of cannula placement for local delivery of MK801 in VMHvl (scale bar = 800 µm).

