## Supplementary figures and images for "Immediate early genes act in the medial hypothalamus to promote adaptation to social defeat"

### Supplementary Figure 1

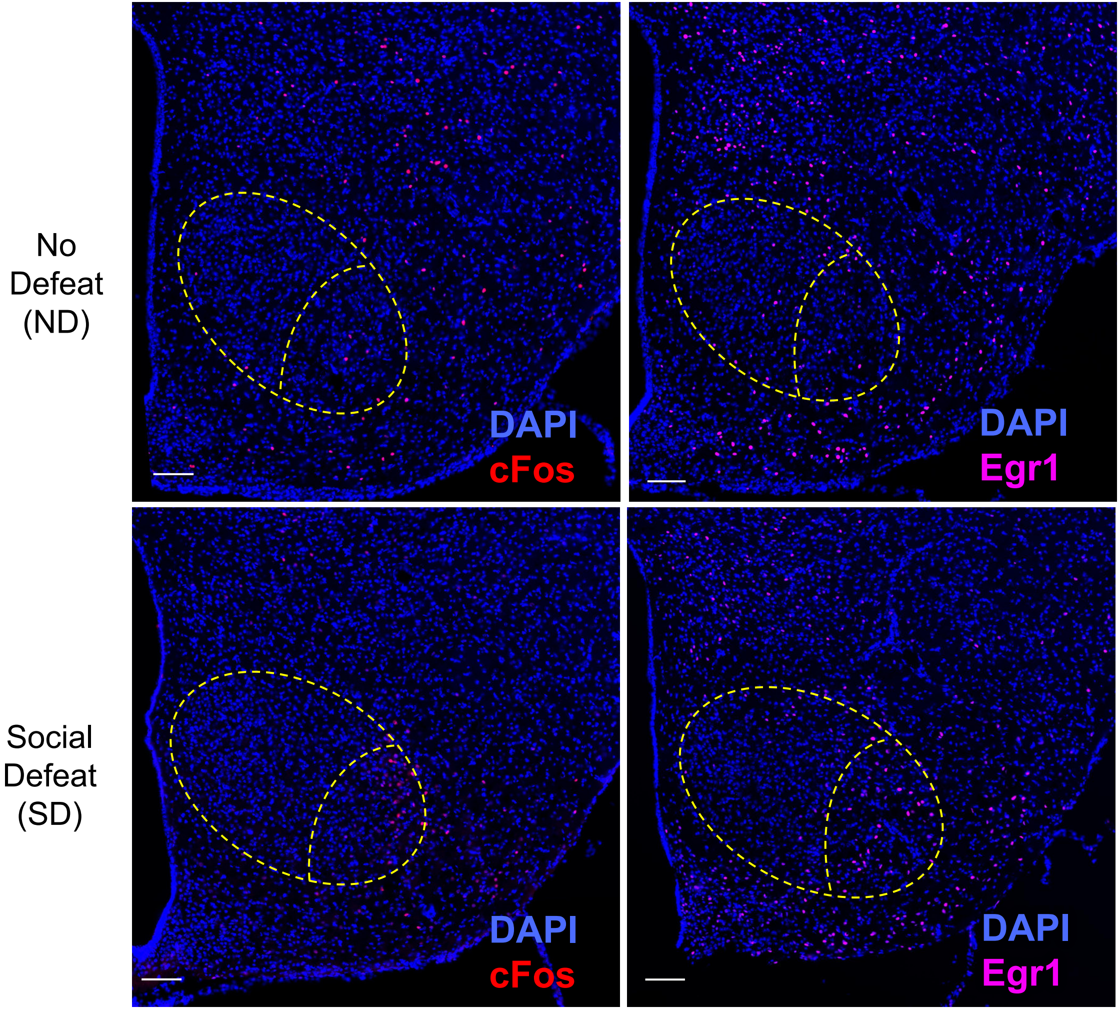

### Supplementary Figure 2

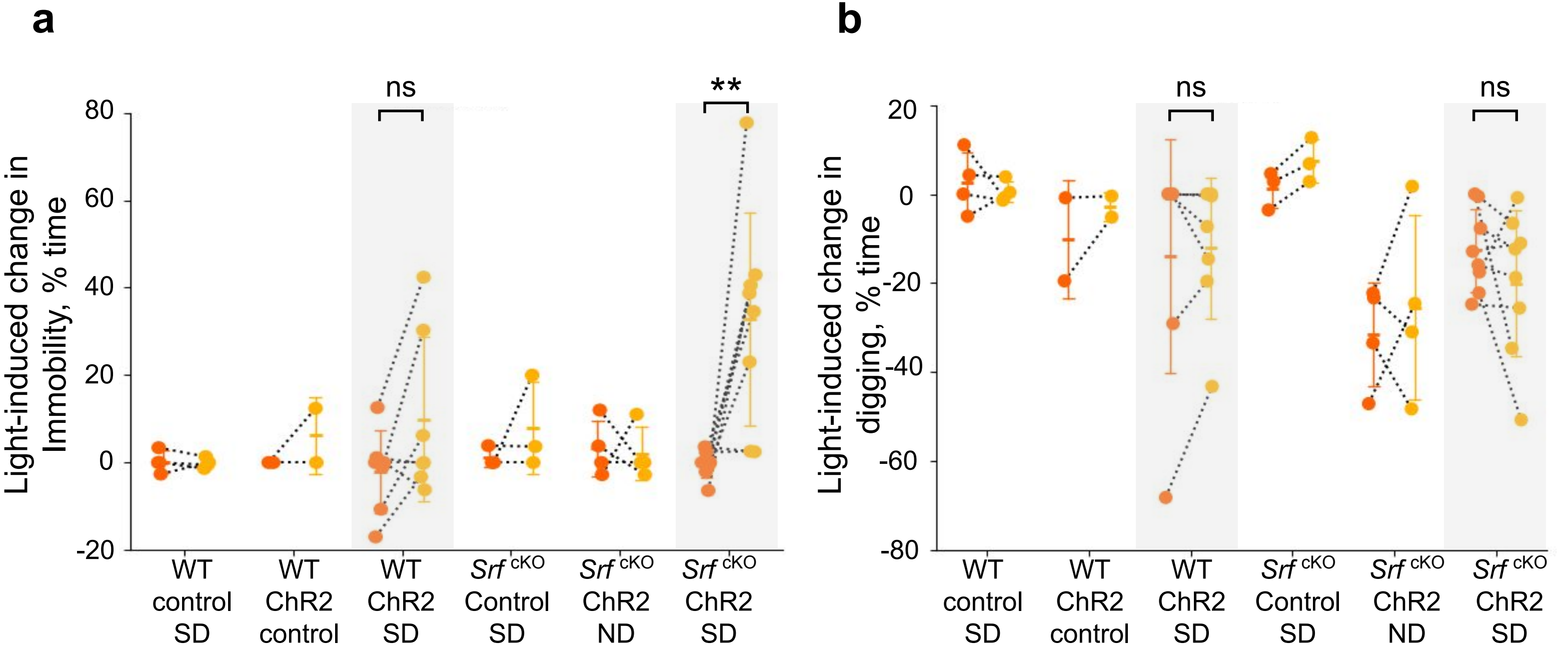

### Supplementary Figure 3

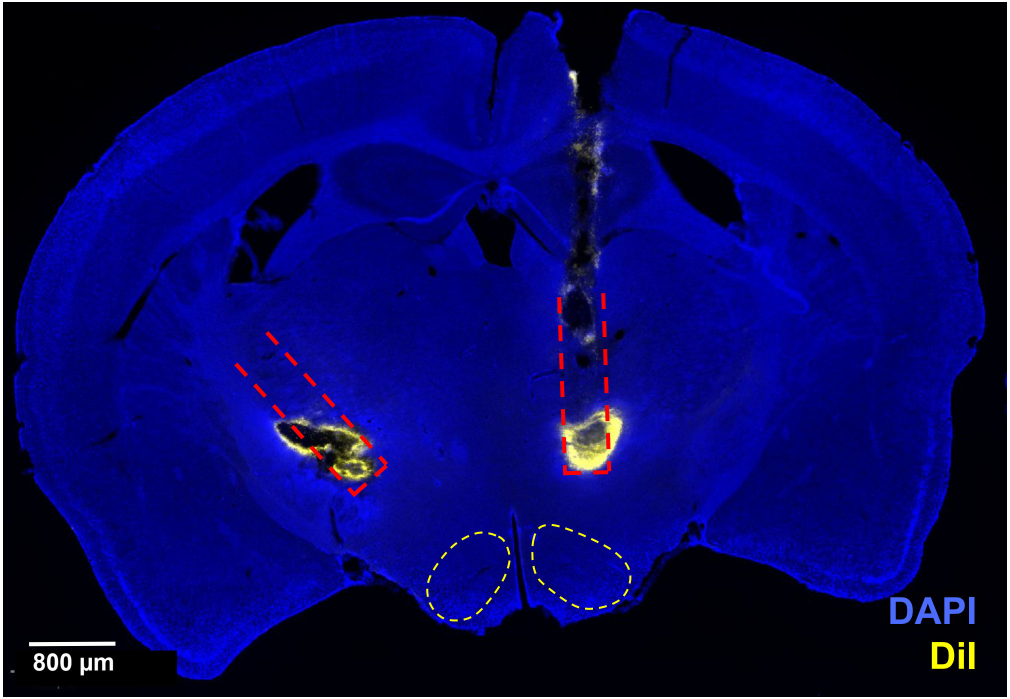
